# Comparison of neural population dynamics in the regression subspace between continuous and categorical task parameters

**DOI:** 10.1101/2022.01.13.476265

**Authors:** He Chen, Jun Kunimatsu, Tomomichi Oya, Yuri Imaizumi, Yukiko Hori, Masayuki Matsumoto, Takafumi Minamimoto, Yuji Naya, Hiroshi Yamada

## Abstract

Neural population dynamics, presumably fundamental computational units in the brain, provide a key framework for understanding information processing in the sensory, cognitive, and motor functions. However, neural population dynamics is not explicitly related to the conventional analytic framework for single-neuron activity, i.e., representational models that analyze neuronal modulations associated with cognitive and motor parameters. In this study, we applied a recently developed state-space analysis to incorporate the representational models into the dynamic model in combination with these parameters. We compared neural population dynamics between continuous and categorical task parameters during two visual recognition tasks, using the datasets originally designed for a single-neuron approach. We successfully extracted neural population dynamics in the regression subspace, which represent modulation dynamics for both continuous and categorical task parameters with reasonable temporal characteristics. Furthermore, we combined the classical optimal-stimulus analysis paradigm for the single-neuron approach (i.e., stimulus identified as maximum neural responses) into the dynamic model, and found that the most prominent modulation dynamics at the lower dimension were derived from these optimal responses. Thus, our approach provides a unified framework for incorporating knowledge acquired with the single-neuron approach into the dynamic model as a standard procedure for describing neural modulation dynamics in the brain.

## Introduction

Recent innovations in the state-space analysis applied to multi-neuronal activities provide insight into the dynamic structure of information processing in a neural population (Brendel et al., 2011; Churchland et al., 2012; Mante et al., 2013). The identified dynamic structures of neural population activity are known as neural population dynamics and are assumed to reflect some underlying computations occurring in a neural network in the sensory, cognitive, and motor domains (Aoi et al., 2020; Churchland et al., 2012; Murray et al., 2017; Okazawa et al., 2021; Osako et al., 2021; Raposo et al., 2014; Rossi-Pool et al., 2021). In the state-space analysis, multi-neuronal interactions with fine temporal evolution have provided a different perspective from the conventional analytical framework for single-neuron activity, known as the representational model. In this conventional framework, the neuronal discharge rate of a single neuron is assumed to reflect some mathematical parameters presumably computed in a neural circuit, such as the Gabor function in the visual cortices (Jones & Palmer, 1987; Tolhurst & Movshon, 1975), movement direction (Georgopoulos et al., 1982) and muscle force (Fetz & Cheney, 1980) in the motor cortices, reward value in the parietal cortex (Platt & Glimcher, 1999), and the location of animals during navigation in the hippocampus (O’Keefe & Dostrovsky, 1971). As dynamic and representational models have rarely been analyzed simultaneously, a fundamental question remains as to how these two different approaches reflect putatively different or shared aspects of neural computation employed by each neuron and the underlying neuronal network, as well as their relationship.

Theoretical neuroscience has provided a quantitative basis for the computation of single neurons in the brain (Dayan & Abbott, 2001). The theory has been developed in parallel with the development of measurement technology for neuronal activity (Yuste, 2015). The early representative model was developed when researchers observed only one neuron while animals performed a behavioral task or were under anesthesia. As the single-neuron recording technique provides fine neuronal activity *in vivo* (Evarts, 1968; Hubel & Wiesel, 1959; Mountcastle & Henneman, 1949; Wurtz, 1968), an analytical and theoretical framework was developed to describe the functional role of separately recorded single-neuron activity. Recently, large-scale multi-channel recording technology has been developed to measure a large number of isolated neurons (Buzsaki et al., 2015; Jun et al., 2017) never imagined before. These simultaneously recorded single-neuron activities in the tens of thousands motivated computational neuroscientists to pursue a theoretical framework for neural computations that provides a different perspective from the conventional representational model (Aoi & Pillow, 2018; Elsayed & Cunningham, 2017; Keemink & Machens, 2019; Saxena & Cunningham, 2019; Vyas et al., 2020).

Neural population dynamics, derived through dimensional reduction of neural population activity and its projection onto parsimonious dimensions, describe the temporal structures of neural response in fine time resolutions in the order of approximately 10 ms, different from other conventional population analyses, e.g., (Georgopoulos et al., 1982). Both analytic frameworks have described brain function in various functional domains, but the relationship between the developing dynamic model and the conventional representational model remains unclear. Indeed, we do not really know whether and how the neural population described by the conventional representational model is described from a dynamic-system perspective. Thus, it is challenging to incorporate knowledge acquired from the representational model into the dynamic model in the form of neural population dynamics.

We previously developed a variant of state-space analysis for continuous parameters (Yamada et al., 2021), which describes how neurons dynamically encode some cognitive parameters in the regression subspace at the population level.

Although the pseudo-population of neurons was composed of non-simultaneously recorded single-neuron activity according to the representational framework, our previous study successfully described neural modulation dynamics using continuous parameters related to value-based decision making. Nevertheless, the other standard parameter for single-neuron recordings, categorical, was not incorporated previously, and thus, our previous analysis were not able to describe all types neural modulations in a dynamic system perspective. Here, we applied our analysis to the pre-existing datasets using a typical factorial design for conventional single-neuron recordings, i.e., categorical task parameters, from the hippocampus in monkeys performing a memory retrieval task (H. Chen & Naya, 2020). Our approach provided the temporal structure of neural modulations for both types of task parameters moment-by-moment, which would not be possible with the representational model, while most aspects of neural modulation were dynamically described, consistent with the conventional representational model. Thus, our analytic approach is beneficial to analyze neural modulation dynamics for all types of pre-existing data allowing researchers to incorporate the representational model into a dynamic system.

## Results

### Task, monkey’s behavior and datasets

Details of the behavioral training, learning progress, and behavioral performance of the animals in the cued lottery task (Exp. 1, Yamada et al., 2021) and in the item– location–retention (ILR) task (Exp. 2, Chen & Naya, 2020) have been previously reported. Briefly, after completing training in Exp. 1, the monkeys learned to estimate the expected value of the lottery, defined as a multiplicative combination of probability and magnitude, and chose the option with higher expected values (Yamada et al., 2021). This choice behavior was observed separately from the neural recordings. We used the neural activity recorded from the central par of the orbitofrontal cortex (cOFC) in the non-choice condition where a single lottery cue and its outcome were provided to the monkeys (Figure 1A–C). In Exp. 2, the monkeys learned to retain the types of visual items and their presented location during the encoding phase, after which the monkeys indicated whether the sample item was matched to the cued items by choosing the memorized location (Figure 1D). Six visual items and four locations were used (Figure 1E). We used the neural activity recorded from the HPC (Figure 1F), after which the sample stimulus was presented to the monkeys during the encoding phase.

**Figure 1.**
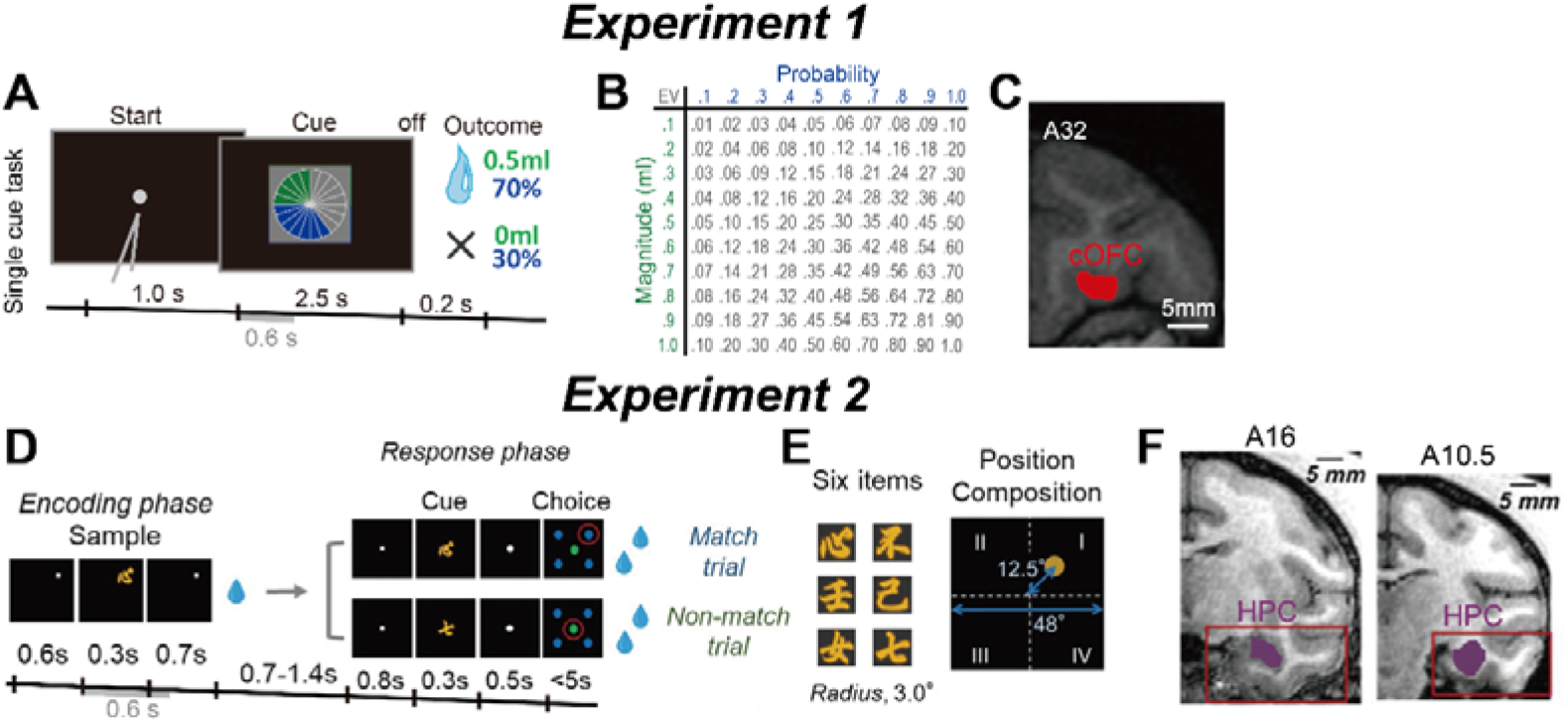
Behavioral task and recording location of neurons. **(A)** Sequence of events during the single-cue task in Exp. 1. A single visual pie chart having green and blue pie segments was presented to the monkeys. Neural activity was analyzed during 0.6 s after cue onset, i.e., for the same duration as in Exp. 2. **(B)** Payoff matrix – each of the magnitudes was fully crossed with each of the probabilities resulting in a pool of 100 lotteries. (**C**) Illustration of neural recording areas based on coronal magnetic resonance (MR) images for the cOFC (13M, medial part of area 13) at the A31–A34 anterior–posterior (A–P) level. (**D**) Sequence of events during the ILR task in Exp. 2. The cue stimulus during the response phase was the same as the sample stimulus during the encoding phase in the match trial, while the two stimuli differed in the nonmatch trial. Neural activity was analyzed during 0.6 s after sample onset, i.e., for the same duration as in Exp. 1. (**E**) Six visual item stimuli and spatial composition during the sample period. (**F**) Coronal MR images from monkey A for the HPC population showing the recording area at A16– A10.5 depicted by purple color in the red boxes. Figure 1A was published in Yamada et al., 2021. Figure 1D–F was published in Chen et al., 2020.

In this study, we constructed two pseudo-simultaneously recorded populations of neurons by aligning the single-neuron activity of the cOFC (Figure 1C, 190 neurons) and HPC (Figure 1F, 590 neurons) with respect to the lottery cue onset in the single-cue task (Figure 1A, gray bar) and the sample onset in the ILR task, respectively (Figure 1D, gray bar), with a 0.6-s time window for each. Note that the HPC population data in Exp. 2 has been analyzed and reported using a representational model, but never analyzed using a dynamic model. Note also that the cOFC population data in Exp. 1 has been analyzed using both representational and dynamic models, and here, we repeated the same analysis with the shorter analysis time window after the cue presentation (2.7 s time window was used in Yamada et al., 2021).

### Conventional analyses for detecting task-dependent modulations

We first applied common conventional analyses such as the general linear model: linear regression in Exp. 1 and ANOVA in Exp. 2, respectively (see Methods). Detailed results from these conventional analyses have been previously reported (Figure 2E–O in Yamada et al., 2021, Figures 2 and 5 in Chen et al., 2020). In Exp. 1, the linear regression analysis showed that the cOFC neurons encode both probability and magnitude to some extent after cue onset, as shown in an example neuron (Figure 2A–B, n = 119 trials, coefficient: intercept, -0.74, t = -0.72, *P* = 0.47; probability, 8.55, t = 6.91, *P* < 0.001; magnitude, 11.1, t = 8.95, *P* < 0.001). This conventional analysis showed whether the probability and magnitude cued by the lottery, both continuous parameters, modulated neuronal activity in each neuron. In the cOFC populations, approximately half of the neurons were modulated by the probability and magnitude of rewards during the 1-s time window (0-1 s after cue onset, probability: 44%, 84/190, magnitude: 49%, 94/190). The analysis with 0.02-s time bins, used to analyze neural population dynamics latter, showed that the percentages of neurons modulated by these two parameters increased and then decreased during the 1.0 s after the onset of the lottery cue (Figure 2C).

**Figure 2.**
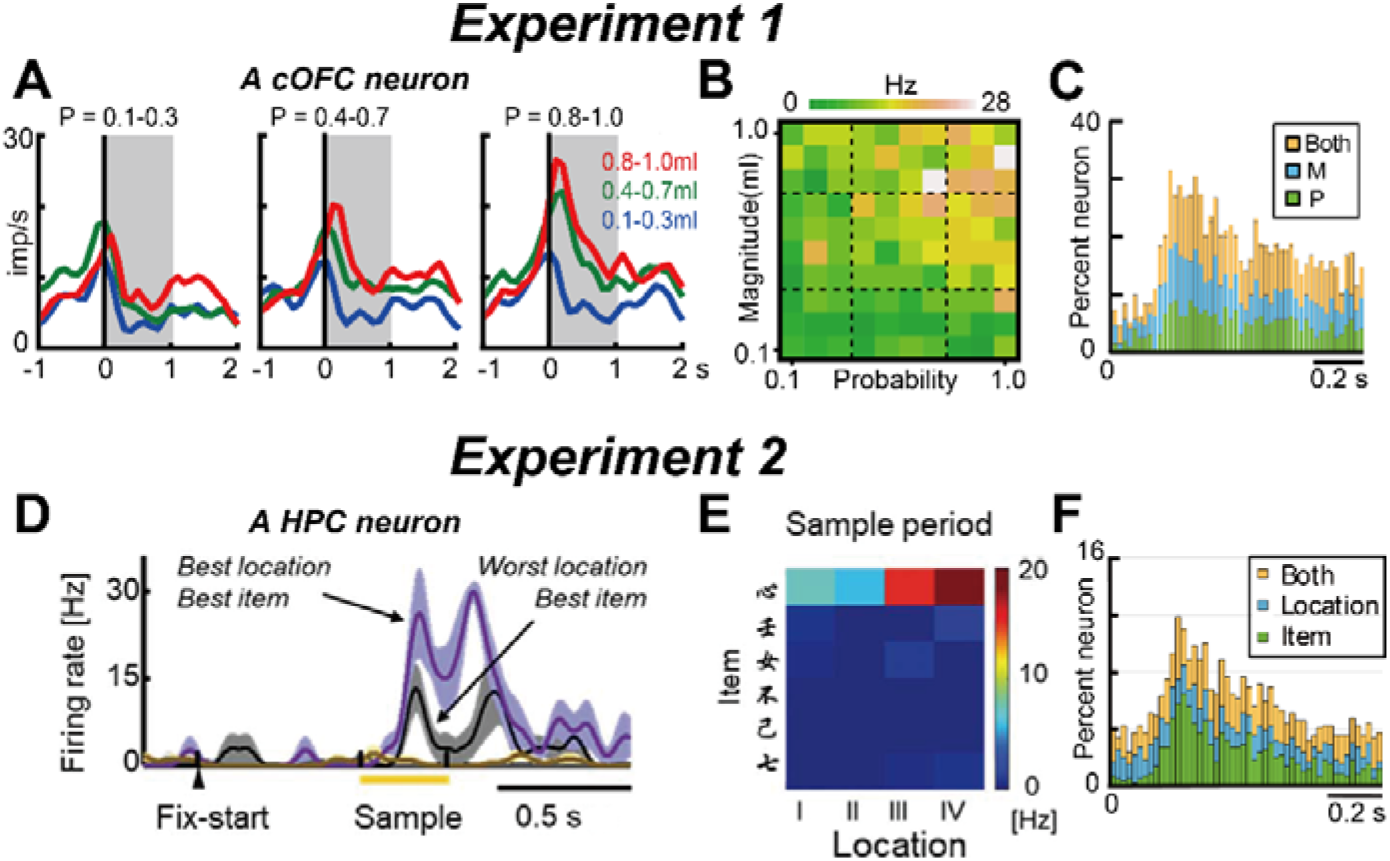
Example activity of neurons during the single-cue and ILR tasks. (**A**) Example activity histogram of a cOFC neuron modulated by the probability and magnitude of rewards during the single-cue task. The activity aligned to the cue onset is represented for three different levels of probability (0.1–0.3, 0.4–0.7, 0.8– 1.0) and magnitude (0.1–0.3 mL, 0.4–0.7 mL, 0.8–1.0 mL) of rewards. Gray hatched time windows indicate the 1-s time window used to estimate the neural firing rates shown in **B**. Histograms are smoothed using a Gaussian kernel ( σ = 50 ms) (**B**) Activity plot of the cOFC neuron during the 1-s time window shown in **A** against the probability and magnitude of rewards. (**C**) Percentages of neural modulation type: the probability (P), magnitude (M), and both (Both) in the 0.02-s time bin during 1.0 s after cue onset. The scale bar indicates the 0.2 s. (**D**) Example of an HPC neuron showing sample-triggered sample–location signals and item signals. A 0.08–0.38 s time window was used to estimate the neural firing rates shown in **E**. Histograms are smoothed using a Gaussian kernel (σ = 20 ms). (**E**) Activity plot of the HPC neuron during the 0.3-s time window shown in **A** against items and locations. (**F**) Percentages of neural modulation types: item, location, and both (Both) in the 0.02-s time bin during 1.0 s after sample onset. Figure 2A–C was published in Yamada et al., 2021. Figure 2D–E was published in Chen et al., 2020.

In Exp. 2, ANOVA showed that the HPC neurons could encode both types of items and their presented locations to some extent, as shown in an example neuron (Figure 2D–E, two-way ANOVA, n = 240 trials, item: F_(5,216)_ = 79.50, *P* < 0.001, location: F_(3,216)_ = 5.48, *P* = 0.001). The analysis showed whether the items and locations, both categorical parameters, modulated neuronal activity in each neuron. In the HPC population, considerable proportions of neurons were modulated by these two factors (0.08–1 s after sample onset, Item, 26%, 152/590, Position, 22%, 131/590). These proportions were significantly smaller than those of the cOFC neurons modulated by the probability and magnitude in Exp. 1 (Chi-squared test, df =1, P < 0.001 for all cases). In the 0.02-s time bins, the percentages of neurons modulated by these two factors increased and then decreased during the 1.0 s after the onset of the sample stimulus (Figure 2F).

In short, the general linear model detected neural modulations using continuous and categorical parameters, which are usually used in the standard representational model, but these analyses did not clearly provide temporal structure of neural population signals.

### State-space analysis for detecting neural modulation dynamics at the population level

State-space analysis originally provided temporal dynamics of neural population signals related to cognitive and motor performances for whole neural activity changes under an assumption of linear system (Churchland et al., 2012; Mante et al., 2013). We previously developed a variant of the state-space analysis, which extracts the temporal structure of neural modulation by the task-related continuous parameters, probability and magnitude of rewards (Yamada et al., 2021). Here, we extend this analysis to neural modulations by categorical parameters to describe how the HPC neural population reflects item and location dynamically. We represented each neural-population signal as a vector time series in the parsimonious dimensions in two steps (Figure 3). First, we used a general linear model to project a time series of each neural activity into a regression subspace composed of task parameters as continuous (Figure 3A) and categorical (Figure 3B) (see Methods for details). This step captures the across-trial variance caused by the task-related parameters moment-by-moment at the population level. Note that this step requires an orthogonal matrix for task parameters because the estimation of the regression subspace is distorted given that the estimation of the regression matrix assumes orthogonality between parameters. Second, we applied PCA once to the time series of neural activities in the regression subspace in each neural population. This step determined the main feature of the neural population signal moment-by-moment in the predominant dimensions at the population level. Because neural activations are dynamic over time, this analysis identified whether and how signal modulations occur as a time-series of eigenvectors. These extracted time series of eigenvectors captured how the main neural modulation evolved as a vector angle and size, and their deviance at the population level (Figure 3C).

**Figure 3.**
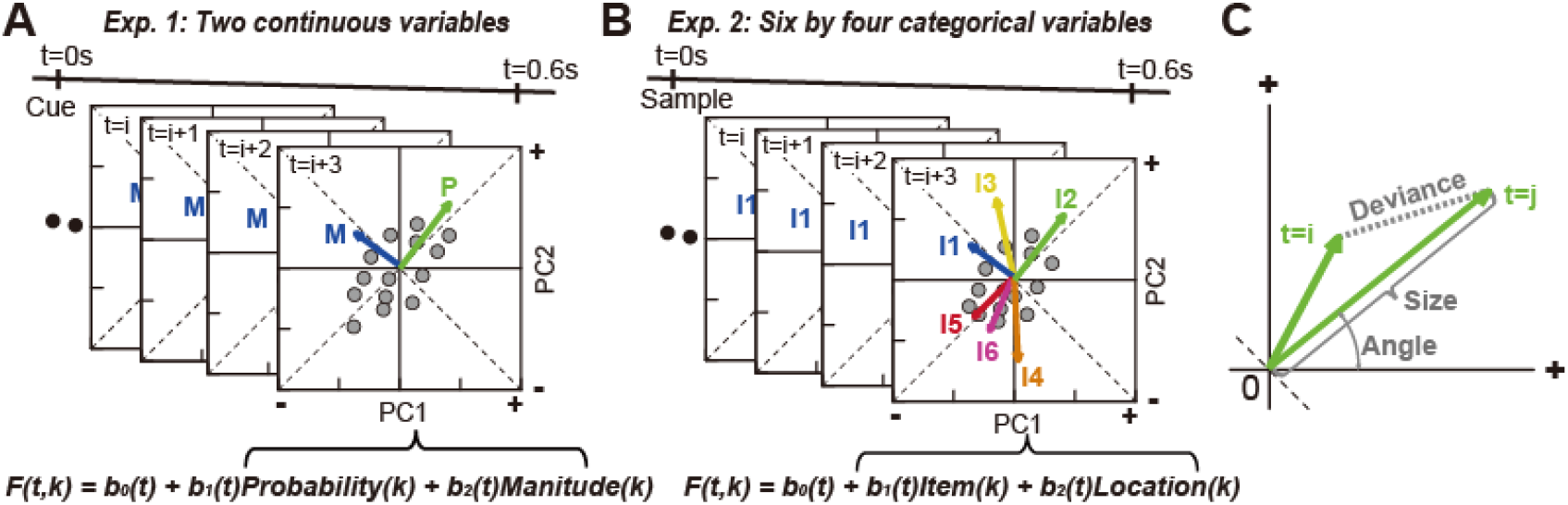
Schematic depictions for the analysis of neural-population dynamics using PCA. (**A**) Time series of neural population activity projected to a regression subspace composed of probability and magnitude. The eigenvectors for probability and magnitude were plotted after coordinate transformation against PC1 and PC2. A series of eigenvectors was obtained by applying PCA once to the cOFC population. The number of eigenvectors obtained by PCA was 0.6 s divided by the analysis window size, 0.02 s, for probability (P) and magnitude (M), hence 30 eigenvectors for each. The regression equation is shown at the bottom (see Methods for details). (**B**) Time series of neural population activity projected to a regression subspace composed of items and locations. The eigenvectors for six items (I1 to I6) were plotted after coordinate transformation against PC1 and PC2 (the eigenvector for locations are not shown). A series of eigenvectors was obtained by applying PCA once to the HPC population. The number of eigenvectors obtained by PCA was 0.6 s divided by the analysis window size, 0.02 s, for the six items and four locations, hence 30 eigenvectors for each. The regression equation is shown at the bottom (see Methods for details). (**C**) Characteristics of the eigenvectors evaluated quantitatively. Angle: vector angle from the horizontal axis obtained from -180^°^ to 180^°^. Size: eigenvector length. Deviance: difference between vectors.

We evaluated the eigenvector properties in the first three principal components (PC1 to PC3) in each neural population in terms of vector angle, size, and deviance. We compared two neural populations recorded during two different cognitive tasks in terms of these vector properties.

### Neural population dynamics reflecting continuous and categorical parameters

Our state-space analysis described the neural population dynamics in the cOFC (Figure 4A–C) during the perception of visual lotteries. In our previous study, we reported neural population dynamics during whole a cue period of 2.7 s (Yamada et al., 2021), but here, we analyzed the dynamics only during the initial 0.6 s to ensure that the neural population structures would be comparable between the cOFC and HPC populations with continuous and categorical parameters. We first confirmed the performance of the state-space analysis indicated by the percentages of variance explained in the cOFC population (Figure 4A). The cOFC population exhibited high performance, more than 40% of the variance was explained by PC1 and PC2 (see gray arrowhead). This is consistent with our previous findings (Figure 7A in Yamada et al., 2021, 27% in 0.02s bin during 2.7 s). We then characterized the whole structure of the cOFC population by plotting its eigenvectors moment-by-moment with the temporal order. As shown in Figure 4B, the eigenvectors for PC1 and PC2 evolved less than 0.2 s after the onset of the cues in both probability and magnitude, while the eigenvectors shortened after approximately 0.3 s. These changes in eigenvectors were very stable in terms of vector angle (Figure 4C, top), as seen in the vector evolutions at 45° in angle between the PC1 and PC2 plane (see also Figure 7B in Yamada et al., 2021), while the vectors changed in PC3 in the opposite direction from positive to negative over time (Figure 4B and Figure 4C, bottom). Thus, these stable structures in the top two dimensions are consistent with our previous ones, even when the analysis window sizes differed.

**Figure 4.**
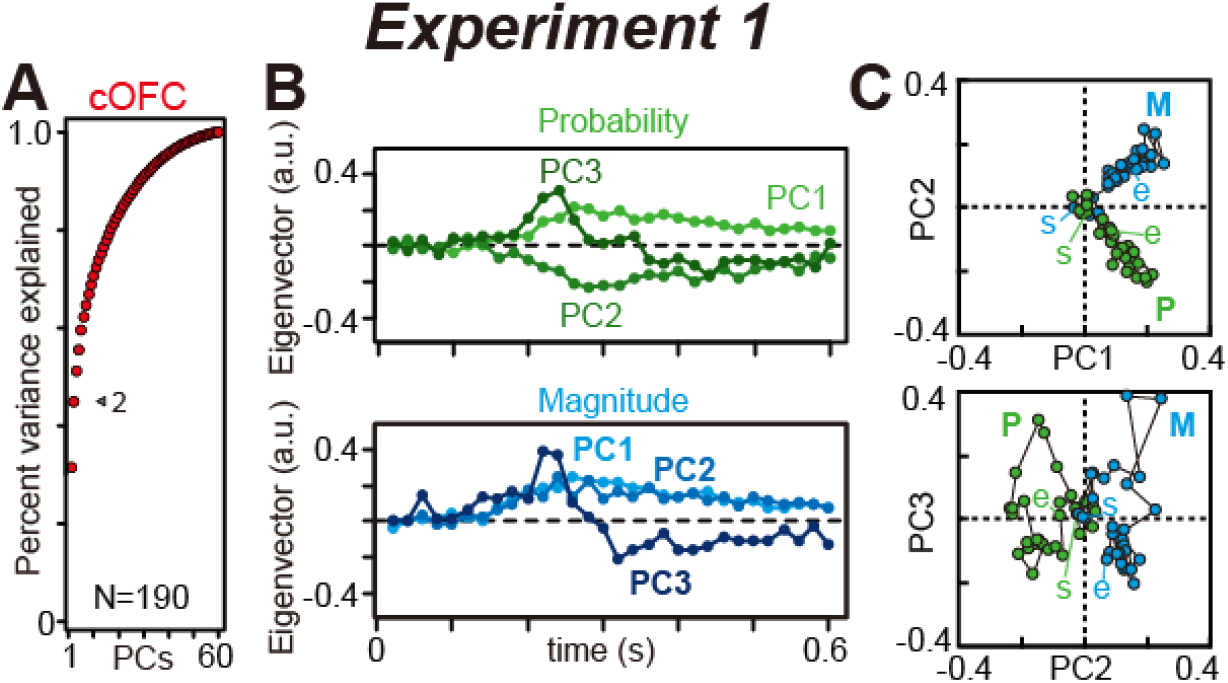
The state-space analysis provides a temporal structure of neural modulation in the cOFC. (**A**) Cumulative variance explained by PCA in the cOFC population. The arrowhead indicates the percentages of variances explained by PC1 and PC2. (**B**) Time series of eigenvectors for PC1 to PC3 in the cOFC population. (**C**) Series of eigenvectors for PC1 to PC3 are plotted against the PC1 and PC2 and PC2 and PC3 dimensions in the cOFC population. Plots at the beginning and end of the series of vectors are labeled as start (s) and end (e), respectively. In A–B, a.u. indicates arbitrary unit.

Upon analysis of the HPC population as modulated by the two categorical parameters, the performance of the analysis was lower than that in Exp. 1 (Figure 5A). The first two PCs only explained approximately 10% of the variance (see gray arrowhead) possibly because the percentages of modulated neurons in the recorded HPC population were not high compared to the cOFC populations (Figure 2C and F). This might also be partly because of the larger data matrix composed of 10 vectors at each time point (six items and four locations) and a larger neural population containing 590 neurons: total *X* of size *N_(590)_*×*M_(300)_* because in our previous study, PCA performance decreased as the matrix size increased (Figure 7A in Yamada et al., 2021). We evaluate the effect of matrix size on PCA performance later in the manuscript (Figure 9). The eigenvectors in the first three PCs appeared to describe the neural population dynamics in the HPC. For example, the extracted eigenvectors for each visual item evolved within a reasonable range of time; increase and then decreased during approximately 0.2 to 0.5 s (Figure 5B), consistent with our previous findings using typical conventional analysis (Figures. 2 and 3 in Chen and Naya, 2020). In clear contrast, the eigenvectors for locations did not show clear trends over time (Figure 5C), as the location information was shown to the monkeys before the sample presentations. When plotting the eigenvectors in the space of the first three PCs, the eigenvectors consistently evolved in one direction in the spaces of PC1 and PC2 (I2, I3, and I6) or in PC3 (I1, I4, and I5) (Figure 5D, left). In contrast, the eigenvectors for the locations were positioned at a constant location across time (Figure 5D, right). Unambiguously, arrangements of the eigenvectors for items and locations were orthogonalized, as seen in the item representations in the second and fourth quadrants and location representations in the first and third quadrants (Figure 5D, top row). Thus, our state-space analysis in the regression subspace successfully described neural modulation dynamics in the HPC populations similar to the cOFC populations, while they reflect continuous and categorical parameters in their neural modulations.

**Figure 5.**
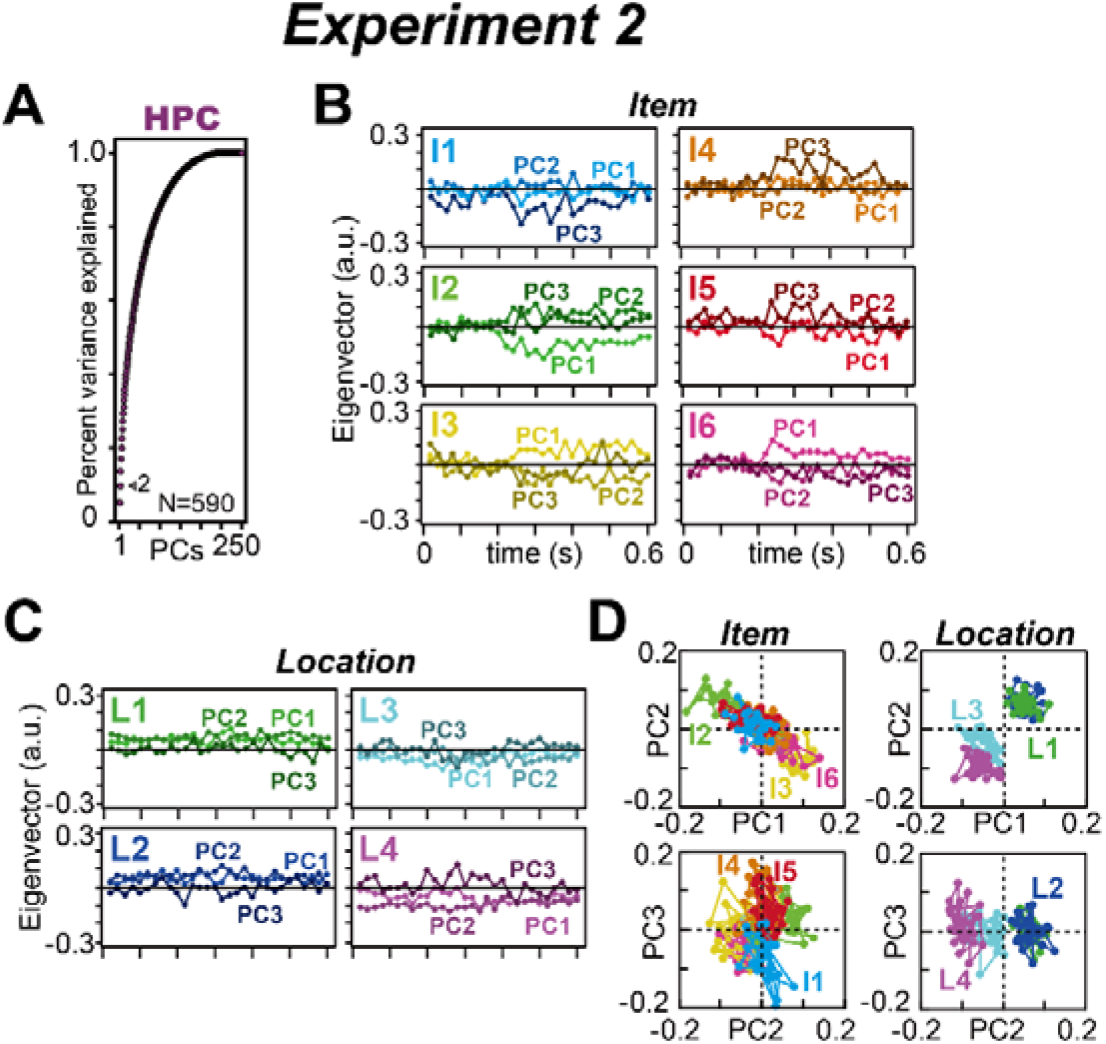
Temporal structure of neural modulation in the HPC population. (**A**) Cumulative variance explained by PCA in the HPC population. The arrowhead indicates the percentages of variances explained by PC1 and PC2. (**B**) Time series of eigenvectors for six items in the HPC population. The top three PCs are shown. (**C**) Same as B but showing the eigenvectors for the four locations. (**D**) Series of eigenvectors for PC1 to PC3 are plotted against the PC1 and PC2 and PC2 and PC3 dimensions in the HPC population. In B–C, a.u. indicates arbitrary unit.

### Effect of shuffle control on PCA performance

To validate the significance of these findings, we used a shuffle control procedure in three ways (see Methods for details), which determines the number of available dimensions in the neural population. In shuffled conditions 1 and 2, information on task-related parameters was partially shuffled in the regression subspace, matrix X. In shuffle condition 1, random permutation of neuron, *n*, was performed at each time *i*, eliminating the temporal neural modulation structure by condition C across each neuron but retaining the effect of neural modulation at each time, *i*, at the population level. In shuffle condition 2, random permutation of time, *i*, was performed in each neuron, *n*, eliminating the temporal neural modulation structure by condition C in each neuron but retaining the effect of neural modulation in each neuron, *n,* at the population level. In shuffled condition 3, random permutation of both time *i* and neuron *n* was performed. We evaluated the performance of the PCAs for each condition of each experiment.

As shown in Figure 6, these three shuffle control procedures reproduced different disturbances in neural populations. In shuffle conditions 1 and 3 (Figure 6A, left and right), the explained variance decreased compared to those from the original data in the cOFC population. In shuffle condition 2, a considerable amount of variance was explained by PCA (Figure 6A, middle). These effects are consistent with those of our previous study (Figure 5A, E, and I in Yamada et al., 2021). Because the eigenvectors were very stable across time in the cOFC population (Figures 4B and 4C), the shuffle within each neuron did not strongly affect PCA performance (Figure 6A, middle). In contrast, the shuffle among neurons at each time point, *t*, strongly reduced the performance of PCA because neural modulation differed neuron-by-neuron (Figure 6A, left). The same effects of shuffle controls were observed in the HPC population, for which categorical parameters were used (Figure 6B); a considerable amount of variance was explained by PCA in shuffled condition 2. When examining the details of the decreased performance in each experiment, the performances of the first three PCs and the first twelve PCs were better than those in shuffled control condition 2 in Exp. 1 and Exp. 2, respectively (P < 0.05 for all these cases). Thus, the total number of available dimensions differed between the experiments. Note that all three shuffles destroyed the structured neural population dynamics to some extent, consistent with our previous findings (Figure 5F and J in Yamada et al., 2021).

**Figure 6.**
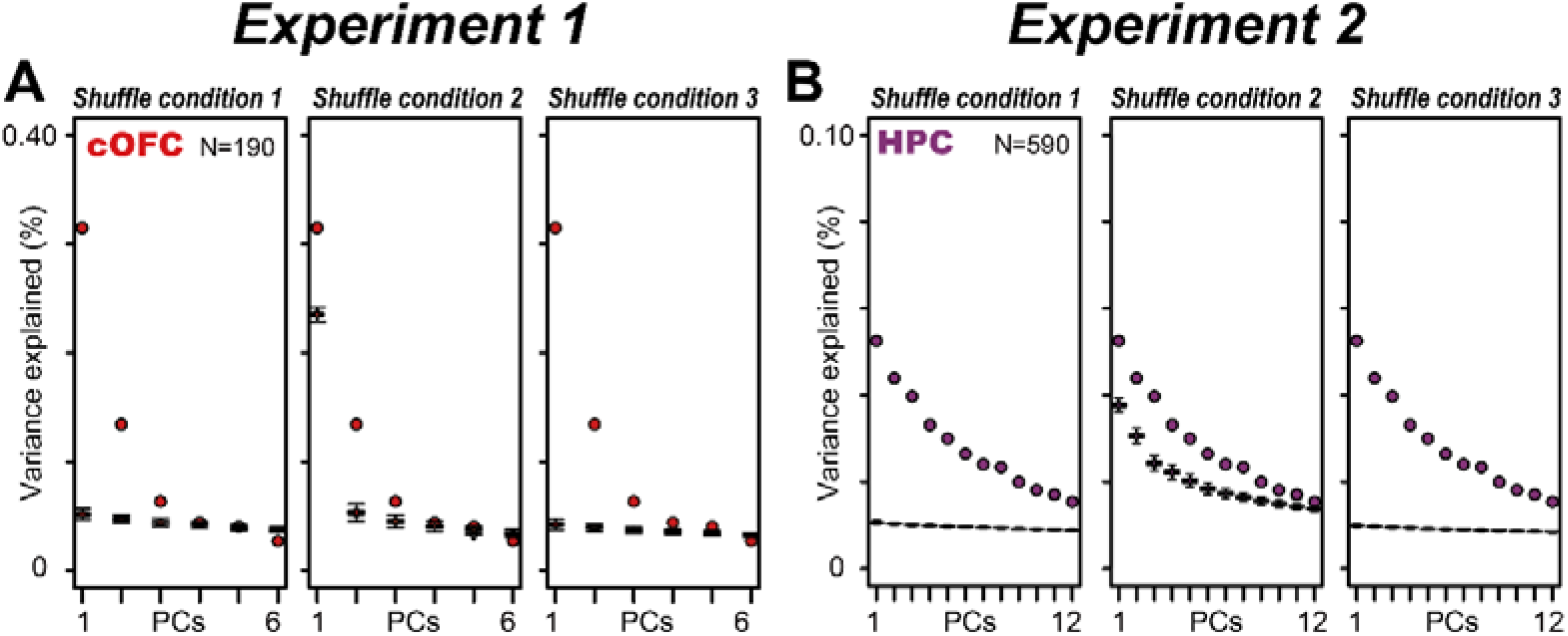
Explained variances by PCA in shuffled controls. (A) Boxplot of explained variances by PCA for PC1 to PC6 for the cOFC population under the three shuffled conditions (see Methods for details). The plot is not cumulative. The boxplot was made with 1,000 repeats of the shuffle in each condition. **(B)** Same as A, but for the HPC population. In **A** and **B**, the colored circles indicate the variances explained by PCA in each neural population without the shuffles.

### Preference ordering in compatible with the representational model

To incorporate the conventional analytic framework into neural population dynamics, we reconstruct the regression subspace in line with the conventional perspective, such as neural preference to task conditions, item and location in this case. We analyzed the most preferred to least preferred conditions for items and locations in each neuron, in which item and location were remapped to the most preferred to least preferred in each condition of item and location neuron-by-neuron, defined using whole activity in the 0.08–0.6 s analysis window in each neuron. Thus, the regression subspace became composed of the same size, total *X* of size *N_(590)_*×*N_(300)_*, but the condition, C, was changed to the most preferred to least preferred items and the most preferred to least preferred locations. The percent variance explained by the model for PC1 and PC2 was almost the same in the preference-ordering analysis (Figure 7A, 11%) compared to the original analysis (Figure 5A, 10%). The composition of the eigenvectors was also similar between the analyses in the PC1 and PC2 dimensions, locating at the second and fourth quadrants from the most preferred (Ib, best item) to the least preferred (Iw, worst item), but they were clearly different in the PC3 dimension, as seen in the most preferred item (Ib) (Figure 7B, left bottom). The composition of eigenvectors for locations was not clearly changed by preference ordering, even for PC3 (Figure 7B, right bottom). Thus, preference ordering may affect the eigenvector compositions at higher dimensions, equal to or more than PC3.

**Figure 7.**
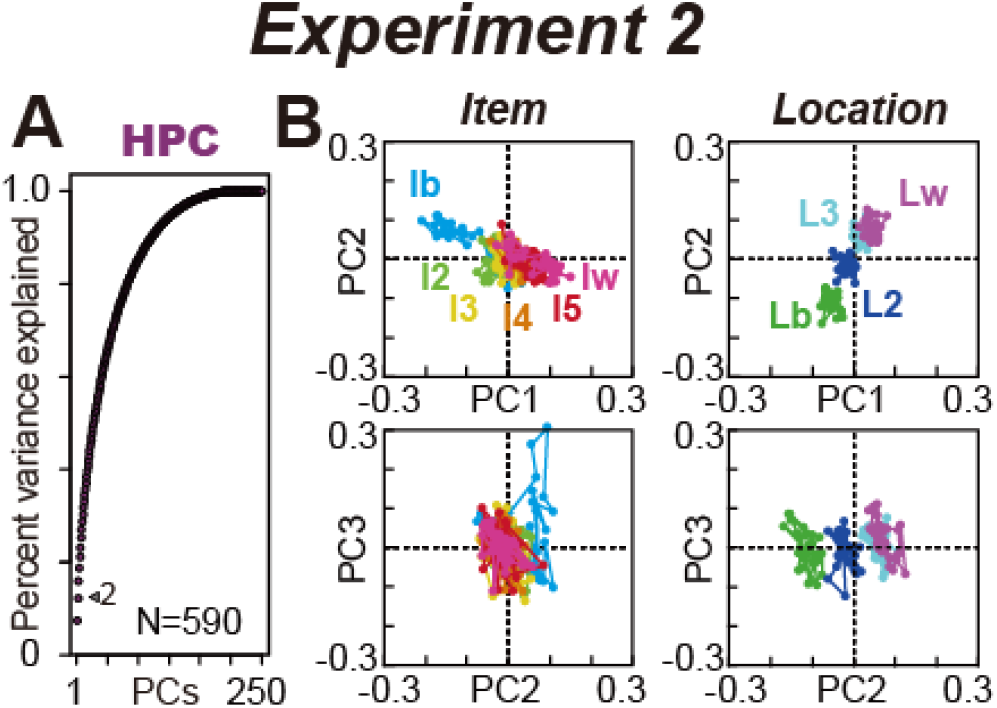
Effects of preference ordering on the HPC categorical data. (**A**) Cumulative variance explained by PCA in the HPC population when item and location are ordered according to their activity preferences (see Methods). The arrowhead indicates the percentages of variances explained by PC1 and PC2. (**B**) Series of eigenvectors for PC1 to PC3 when item and location are ordered according to their preferences plotted against the PC1 and PC2 and PC2 and PC3 dimensions in the HPC population. Ib and Iw indicate the best and worst items, respectively. I2 to I5 indicate the 2^nd^ to 5^th^ best items. Lb and Lw indicate the best and worst locations, respectively. L2 and L3 indicate the 2^nd^ and 3^rd^ best locations, respectively.

### Quantitative analyses of neural population dynamics between two neural populations

To quantitatively examine these neural population structures, we compared the properties of the eigenvectors by estimating the vector size, angle, and deviance in each neural population (Figure 8). For this analysis, we used the rank-ordered HPC data shown in Figure 7, as well as the cOFC data shown in Figure 4. In the rank-ordered data, we evaluated the best and worst conditions as typically used in conventional representational analyses.

**Figure 8.**
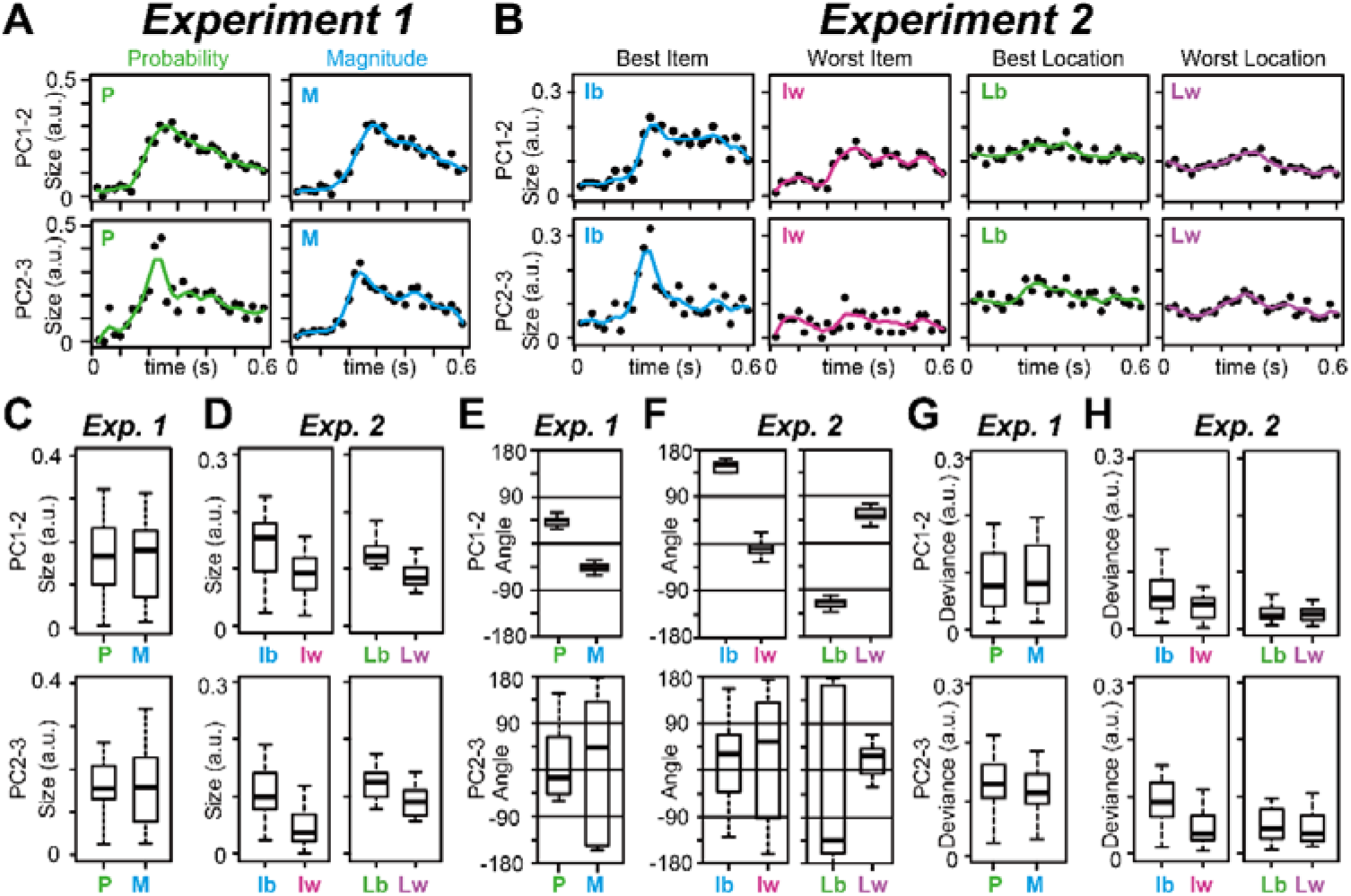
Quantitative evaluations of eigenvector properties in the cOFC and HPC populations. (**A**) Time series of vector size estimated in the cOFC population for probability (P) and magnitude (M) of rewards. The vector sizes are estimated in the PC1 to PC2 plane (top) and PC2 to PC3 plane (bottom), respectively. a.u. indicates arbitrary unit. The solid colored lines indicate interpolated lines using a cubic spline function to provide a resolution of 0.005 s. (**B**) Same as A, but for the best and worst items and the best and worst locations in the HPC population. (**C**) Box plots of vector size estimated in the cOFC population for probability and magnitude of rewards. (**D**) Same as C, but for the best and worst items and the best and worst locations in the HPC population. (**E–F**) Same as C–D, but for the vector angle estimated in the cOFC and HPC populations. (**G–H**) Same as C–D, but from the vector deviance for the mean estimated in the cOFC and HPC populations. In C–H, data after 0.1 s are used.

First, evaluation of vector size provided clear time-dependent structures in both cOFC and HPC populations for probability and magnitude (Figure 8A) and for the best and worst items (Figure 8B). Such time-dependent changes were not clearly observed in the eigenvectors for the best and worst locations (Figure 8B, right and second right columns), presumably because location information had already been provided to the monkeys before the samples appeared. The vector sizes during 0.1 s to 0.6 s after the onset of the lottery stimuli were not significantly different between two continuous parameters, probability and magnitude of rewards (Figure 8C, Wilcoxon rank sum test; PC1-2, n = 52, df = 51, W = 330, *P* = 0.892, PC2-3, n = 52, df = 51, W = 341, *P* = 0.964), consistent with our previous findings (Yamada et al., 2021). In contrast, the vector sizes during 0.1 s to 0.6 s after the onset of the sample stimuli significantly differed between the best and worst items (Figure 8D, Wilcoxon singed rank test; PC1-2, item, n = 52, df = 51, W = 502, *P* = 0.002, PC2-3, item, n = 52, df = 51, W = 588, *P* < 0.001; PC1-2, location, n = 52, df = 51, W = 600, *P* < 0.001, PC2-3, item, n = 52, df = 51, W = 542, *P* < 0.001), possibly because the regression coefficients for the best conditions were considerably different from their means because the HPC responses were highly selective for one object (Figure 2D–E). Thus, the vector sizes captured the temporal changes in neural modulation at the population level.

The analyses of vector angles showed that all eigenvectors were very stable in both populations in the top two dimensions (Figure 8E–F, top, Wilcoxon rank sum test; cOFC, PC1-2, n = 52, df = 51, W = 62, *P* < 0.001; HPC, PC1-2, item, n = 52, df = 51, W = 520, *P* < 0.001, location, n = 52, df = 51, W = 0, *P* < 0.001), as also shown in Figures 4C top and 7B top. Their angles in the PC2-3 plane were not stable (Figure 8E–F, bottom, Wilcoxon rank sum test; cOFC, PC2-3, n = 52, df = 51, W = 343, *P* = 0.935; HPC, PC2-3, item, n = 52, df = 51, W = 321, *P* = 0.765, PC2-3, location, n = 52, df = 51, W = 312, *P* = 0.643, see also, Figure 4C, bottom and Figure 7B, bottom). Both neural populations showed considerable vector deviance smaller than 0.1 with some statistical differences (Figure 8G–H, Wilcoxon rank sum test; cOFC, PC1-2, n = 52, df = 51, W = 361, *P* = 0.683; PC2-3, n = 52, df = 51, W = 300, *P* = 0.496; HPC, PC1-2, item, n = 52, df = 51, W = 459, *P* = 0.027; PC2-3, item, n = 52, df = 51, W = 581, *P* < 0.001, PC1-2, location, n = 52, df = 51, W = 352, *P* = 0.807; PC2-3, location, n = 52, df = 51, W = 384, *P* = 0.408). Thus, our state-space analysis in the regression subspace was capable of describing neural modulation dynamic in the cOFC and HPC during two different cognitive tasks composed of continuous and categorical parameters.

### Matrix size control for PCA

Because the PCA performance was lower in the HPC than in the cOFC population, we evaluated the effect of matrix size on the representational models. In our previous study, the variance explained by the PCA decreased as the matrix size increased to explain the same neural modulation (Figure 7A in Yamada et al., 2021). In the present study, we reduced the matrix size of the HPC population by extracting the best and worst conditions for item and location according to conventional representational model analysis, although the regression matrix from the other conditions, 2^nd^ preferred to 5^th^ preferred, were removed. The regression subspace was reduced from the large size, total *X* of size *N_(590)_*×*N_(10×30)_*, to *N_(590)_*×*N_(4×30)_,* similar column size to the OFC population, *N_(190)_*×*N_(2×30)_*, in terms of the number of conditions. In this smaller regression matrix, PCA performance improved (Figure 9A, approximately 16% of the variance explained by PC1 and PC2), consistent with the findings of our previous study where we used continuous parameters. The eigenvector compositions developed in a clearly symmetric way, perhaps because the variances from the other conditions were removed (Figure 9B). In this smaller regression matrix, the principal components appeared to be rotated at an approximately 135° angle from the original on the PC1-2 plane (Figures 9B and 7B).

The percent variance explained by the PCA clearly differed from that in the shuffled conditions for the top three PCs, while the top six PCs significantly differed from shuffled control in condition 2 (Figure 9C, see also Figure 6B, middle), indicating that some neural population structures in higher dimensions were removed in this smaller matrix.

**Figure 9.**
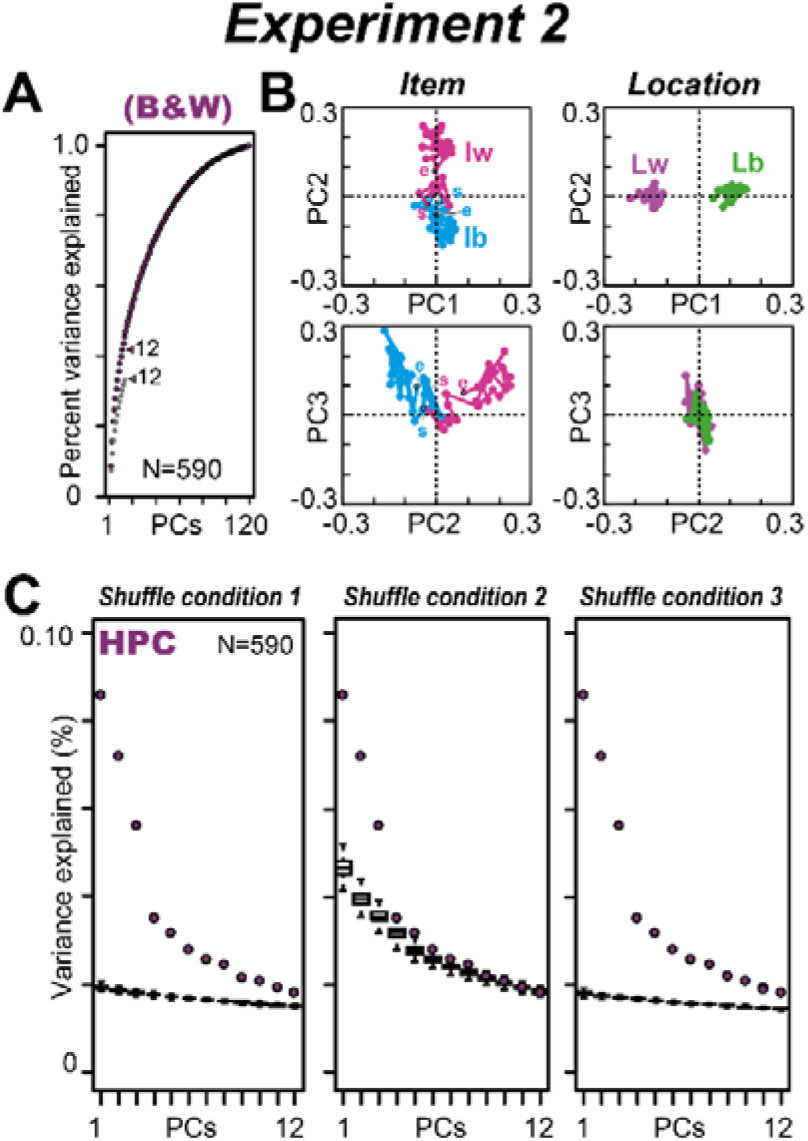
Effects of matrix size control in the HPC population. (**A**) Cumulative variance explained by PCA in the HPC population when the best and worst conditions for item and location are used for the regression subspace. The gray dots indicated the percent variance explained by the PCA when using the full matrix. The first 12 PCs are shown. (**B**) Time series of eigenvectors for PC1 to PC3 when the best and worst items and the best and worst locations are used. Ib and Iw indicate the best and worst items, respectively. Lb and Lw indicate the best and worst locations, respectively. s and e indicate the start and end of the time series of vectors, respectively. (**C**) Boxplot of explained variances by PCA for PC1 to PC12 under the three shuffled conditions (see Methods for details). The plot is not cumulative. The boxplot was made with 1,000 repeats of the shuffle in each condition. The colored circles indicate the variances explained by PCA in the HPC population without the shuffles.

In summary, our state-space analysis clearly described the neural modulation structures for both continuous and categorical task parameters. In both populations, using two standard task designs, we found stable evolutions of neural modulation structures in a relatively short period i.e., 0.6 s while the monkeys perceived visual items.

## Discussion

In our previous study, we developed a variant of state-space analysis in the regression subspace for continuous task parameters, which extracts neural modulation dynamics at the population level. Here, we applied our state-space analysis in the regression subspace to categorical task parameters and successfully described the neural modulation dynamics for items and locations for the first time (Figure 7). Comparisons of these results with those derived from continuous task parameters (Figures 4 and 5) indicated that our analysis showed gradual development (Figure 8A–B) and stable composition of the neural population structures at different angles (Figure 8E–F, top). Moreover, the population analysis using the best and worst conditions for items and locations showed that low-dimensional robust neural-modulation structures existed in this restricted neural population, and some high-dimensional information seemed to disappear by removing neural activity between the best and worst conditions (Figure 9A, and Figures 7B vs 9B). Although both the cOFC and HPC neural populations were pseudo-populations of neurons using repetitive single-neuron recordings for the representational models, we successfully extracted both neural modulation dynamics with the state-space analysis we developed. Our reliable extraction of neural modulation dynamics indicated that any type of data can be re-analyzed and evaluated to describe the temporal structure of neural modulations as dynamic representational models.

### Two different types of task parameters yield comparable regression subspaces

In our state-space analysis, neural population activity was projected to the regression subspace, reflecting the across-trial variance caused by the task-related parameters at the population level. In this step, both continuous and categorical task parameters are reliably used within a framework in the general linear model. However, it was reliably performed with one critical limitation; the conditions in any parameters should be orthogonalized as the experimental design (Grafen & Hails, 2002). In the linear system assumed here, the concept of orthogonality is critical in terms of statistics and to avoid the skewed projection of neural activity into the regression subspace, which is part of the whole neural activity reflecting activity modulation by the task parameters of interest.

Analysis of the regression subspace has been performed in a limited number of studies (Aoi et al., 2020; Mante et al., 2013). These studies aimed at detecting the regression subspace within a whole neural structure at a constant time point (Mante et al., 2013), and the detected modulation axis is assumed to be projected orthogonally and sometimes being stable through a task trial (Aoi et al., 2020). Our results support these assumptions, which were not examined in the previous study, as the cOFC and HPC showed stable evolution of these neural-modulation structures, at least during the two cognitive tasks with continuous and categorical task parameters. Thus, our approach encourages research that combines the conventional representational model and the dynamic model by re-analyzing the pseudo-population of recorded single-neuron activity to remap the dynamic neural modulation structures for all pre-existing data.

### Stable and fluctuating signals in neural modulation dynamics

In this study, we observed stable neural modulation dynamics in both the cOFC and HPC populations. Although these tasks were designed with different types of task parameters, both brain regions showed stable modulation structures during the visual perception (Figures 4, 7, and 8). Why do these two distinct brain regions show stable modulation dynamics? One possibility is that both the cOFC and HPC play a role in accessing the memory for the expected values as a combination of probability and magnitude in Exp. 1 and the association between stimulus and position for future decisions in Exp. 2. These types of stable structures were observed in the dorsolateral prefrontal cortex during a typical working memory task (Murray et al., 2017). Thus, a key aspect of stable neural dynamics may be continuous access to memory and its maintenance.

In our previous study, fluctuating neural population signals were observed in the dorsal part of the striatum (DS) and medial part of the orbitofrontal cortex (mOFC) because of signal instability or weakness (Figure 5A and B in Yamada et al., 2021). Because the signal carried by the mOFC population was weak (Figure 8 bottom row in Yamada et al., 2021), the eigenvector fluctuation in the mOFC population reflected weak signal modulations by the probability and magnitude of rewards. In this case, moment-by-moment vector fluctuation was observed, as there was no clear neural modulation structure in the mOFC populations. In contrast, the fluctuating DS signal seemed to reflect the functional role employed by the DS neural population in detecting and integrating the probability and magnitude of rewards, related to the control of some actions (Balleine et al., 2007). In the DS population, structural changes in eigenvectors occurred over time (Figure 8 in Yamada et al., 2021). We need to elaborate on the stability of modulation dynamic functions in neural processing in future studies to elucidate how neural circuitry actually operates and computes (Ebitz & Hayden, 2021; Humphries, 2021).

## Conclusions

Representational models have provided mounting evidence that neural modulation is associated with mathematical functions in every area of the brain. A dynamic-model approach that has been recently developed appears promising to account for different aspects of neural computation, but the relationship with the representational models remains unclear. Although a few studies have sought a connection between these two advances (X. Chen & Stuphorn, 2015; Churchland et al., 2012; Murray et al., 2017), more direct comparisons are necessary to understand the functional significance of the neural population dynamics. Our results indicated that the neural modulation dynamics observed in population ensemble activities are compatible with representational models and encourage research aimed at incorporating traditional representational models into the dynamic system.

## Materials and Methods

### Subjects and experimental procedures

Four macaque monkeys were employed for this study in two experiments (Experiment 1: *Macaca mulatta,* SUN, 7.1 kg, male; *Macaca fuscata*, FU, 6.7 kg, female; Experiment 2: *Macaca mulatta,* A, 9.3 kg, male; *Macaca mulatta,* D, 9.5 kg, male). All experimental procedures were approved by the Animal Care and Use Committee of the University of Tsukuba (Exp. 1, protocol no H30.336), and the Institutional Animal Care and Use of Laboratory Animals approved by Peking University (Exp. 2, project number Psych-YujiNaya-1) and performed in compliance with the US Public Health Service’s Guide for the Care and Use of Laboratory Animals.

### Behavioral task and Monkey electrophysiology

#### Experiment 1

##### Cued lottery tasks

Animals performed one of two visually cued lottery tasks: *a single-cue task* or *a choice task*. Neuronal activity was recorded only during the single-cue task.

At the beginning of trials during the single-cue task, the monkeys had 2 s to align their gaze to within 3^°^ of a 1^°^-diameter gray central fixation target. After fixation for 1 s, a pie chart was presented for 2.5 s to provide information regarding the probability and magnitude of rewards at the same location as the central fixation target. The probability and magnitude of rewards were associated with the number of blue and green 8^°^ segments, ranging from 0.1 to 1.0 mL in 0.1-mL increments for magnitude and 0.1 to 1.0 in 0.1 increments for probability. With an interval of 0.2 s after the removal of the pie chart, either a 1 kHz or 0.1 kHz tone of 0.15-s duration was provided to indicate reward or no-reward outcomes, respectively. With an interval of 0.2 s after the high tone, a fluid reward was delivered. After a low tone, no reward was delivered. An inter-trial interval of 4–6 s followed each trial.

In the trials during choice task, the animals were instructed to choose between two peripheral pie charts providing information regarding the probability and magnitude of rewards for each of the two target options were presented for 2.5 s, at 8^°^ to the left and right of the central fixation location. The animals received a fluid reward, indicated by the green pie chart of the chosen target, with the probability indicated by the blue pie chart; otherwise, no reward was delivered.

One hundred pie charts were used in the experiments. In the single-cue task, each pie chart was presented once in a random order. In the choice task, two pie charts from the 100 pie charts were randomly allocated to the two options. During one session of electrophysiological recording, approximately 30 to 60 trial blocks of the choice task were interleaved with 100 to 120 trial blocks of the single-cue task.

We used conventional techniques for recording single-neuron activity from the central part of the orbitofrontal cortex (cOFC, area 13M). A tungsten microelectrode (1–3 MΩ, FHC) was used to record single-neuron activity. Electrophysiological signals were amplified, band-pass filtered (at 50–3000 Hz), and monitored. Single-neuron activity was isolated based on the spike waveforms. We recorded from the cOFC of a single hemisphere in each of the two monkeys: 190 cOFC neurons (98, SUN and 92, FU). The activity of all single neurons was sampled when the activity of an isolated neuron demonstrated a good signal-to-noise ratio (>2.5). Blinding was not performed. The sample sizes required to detect effect sizes (number of recorded neurons, number of recorded trials in a single neuron, and number of monkeys) were estimated based on previous studies (X. Chen & Stuphorn, 2015; Yamada et al., 2013; Yamada et al., 2018). Neural activity was recorded during 100–120 trials of the single-cue task. Neural activity was not recorded during the choice trials. In this study, we analyzed the cOFC activity data during 600 ms after cue onset from Yamada et al. (2021) for comparison with the activity data in Exp. 2.

#### Experiment 2

##### Item location-retention (ILR) task

The animals performed the task under dim light in an electromagnetically shielded room. The task started with an encoding phase, which was initiated by the animal pulling a lever and fixating on a white square (0.6°) presented within one of the four quadrants at 12.5° (monkey A) or 10° (monkey D) from the center of the touch screen (3M^TM^ MicroTouch^TM^ Display M1700SS, 17 inch), situated approximately 28 cm from the subjects. Eye position was monitored using an infrared digital camera with a sampling frequency of 120 Hz (ETL-200, ISCAN). After fixation for 0.6 s, one of the six items (3.0° for monkey A and 2.5° for monkey D, radius) was presented in the same quadrant as a sample stimulus for 0.3 s, followed by another 0.7-s fixation on the white square. If the fixation was successfully maintained (typically, < 2.5 °), the encoding phase ended with the presentation of a single drop of water.

The encoding phase was followed by a blank interphase delay interval of 0.7–1.4 s during which no fixation was required. The response phase was initiated with a fixation dot presented at the center of the screen. One of six items was then presented at the center for 0.3 s as a cue stimulus. After another 0.5-s delay period, five disks were presented as choices, including a blue disk in each quadrant and a green disk at the center. When the cue stimulus was the same as the sample stimulus, the animal was required to choose by touching the blue disk in the same quadrant as the sample (i.e., match condition). Otherwise, the subject was required to choose the green disk (i.e., non-match condition). If the animal made a correct choice, four to eight drops of water were provided as a reward; otherwise, an additional 4 s was added to the standard intertrial interval (1.5–3 s). During the trial, a large gray square (48° on each side, Red, Green, Blue value: 50, 50, 50, luminance: 3.36 cd/m^2^) was presented at the center of the display (backlight luminance: 0.22 cd/m^2^) as a background. After the end of the trial, all stimuli disappeared, and the entire screen displayed a light red color during the intertrial interval. The start of a new trial was indicated by the reappearance of the large gray square on the display, upon which the monkey could start pulling the lever, triggering the appearance of a white fixation dot. In the match condition, sample stimuli were pseudo-randomly chosen from six well-learned visual items, and each item was presented pseudo-randomly within the four quadrants, resulting in 24 (6 × 4) configuration patterns. In the nonmatch condition, the location of the sample stimulus was randomly chosen from the four quadrants, and the cue stimulus was randomly chosen from the five items that differed from the sample stimulus. The match and non-match conditions were randomly presented at a ratio of 4:1, resulting in 30 (24 + 6) configuration patterns. The same six stimuli were used during all recording sessions.

To record single-unit activity, we used a 16-channel vector array microprobe (V1 X 16-Edge, NeuroNexus), a 16-channel U-Probe (Plexon), a tungsten tetrode probe (Thomas RECORDING), or a single-wire tungsten microelectrode (Alpha Omega). We recorded 590 hippocampal (HPC) neurons, of which the recording sites appeared to cover all its subdivisions (i.e., dentate gyrus, CA3, CA1, and subicular complex). We applied state-space analysis to the HPC population and compared to the results from the cOFC population.

### Statistical analysis

For statistical analysis, we used the statistical software package R (Exp. 1) and MATLAB (MathWorks) (Exp. 2). All statistical tests for the neural analyses were two tailed.

### Behavioral analysis

Exp. 1. We previously reported that monkey behavior depends on the expected values defined as probability time magnitude (Yamada et al 2021).

Exp. 2. We previously reported that two monkeys learned to retain the item and location information of the sample stimulus (H. Chen & Naya, 2020).

No new behavioral results were included in this study.

### Neural analysis

Peristimulus time histograms were drawn for each single-neuron activity aligned at the visual stimulus onset. The average activity curves were smoothed for visual inspection using a Gaussian kernel.

### Conventional analyses to detect neural modulations in each neuron

We analyzed neural activity during the 1-s time window (0-1 s after cue onset, Exp. 1) and during the 0.92 s time window (0.08-1 s after sample onset, Exp. 2), respectively, respectively. These activities were used for the conventional analyses below. No Gaussian kernel was used.

*Exp. 1.* Neural discharge rates (*F*) were fitted using a linear combination of the following parameters:

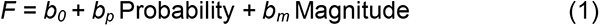

where Probability and Magnitude are the probability and magnitude of the rewards indicated by the pie chart, respectively. *b*_0_ is the intercept. If *b_p_* and *b_m_* were not 0 at *P* < 0.05, the discharge rates were regarded as being significantly modulated by that variable. These results have been previously reported (Yamada et al., 2021).

Based on the linear regression, activity modulation patterns were categorized into several types: “Probability” type with a significant *b*_p_ and without a significant *b*_m_; “Magnitude” type without a significant *b*_p_ and with a significant *b*_m_; “Both” type with significant *b*_p_ and *b*_m_.

*Exp. 2.* For neural responses during the encoding phase after the sample presentation, we evaluated the effects of “item” and “location” for each neuron using two-way analysis of variance (ANOVA) (*P* < 0.01 for each). We analyzed neurons that we tested in at least 60 trials (10 trials for each stimulus, 15 trials for each location). On average, we tested 100 trials for each neuron (n = 590). The results have been previously reported (H. Chen & Naya, 2020).

Based on the ANOVA, activity modulation patterns were categorized into several types: “Item” type only with a significant main effect of Item; “Location” type only with a significant effect of Location; “Both” type with a significant effect of Item and Location or with a significant effect of interaction.

### Population dynamics using principal component analysis

We analyzed neural activity during a 0.6 s time period from cue onset (Exp. 1) and sample onset (Exp. 2). To obtain a time series of neural firing rates within this period, we estimated the firing rates of each neuron for every 0.02-s time bin (without overlap) during the 0.6-s period. No Gaussian kernel was used.

#### Regression subspace

We used a general linear model to determine the probability and magnitude of rewards (Exp. 1) and item and location (Exp. 2) affecting the activity of each neuron in the neural populations. Each neural population was composed of all recorded neurons in each brain region.

*Exp. 1.* We first set the probability and magnitude at 0.1 and 1.0 and 0.1 to 1.0 mL, respectively. We then described the average firing rates of neuron *i* at time *t* as a linear combination of the probability and magnitude in each neural population:

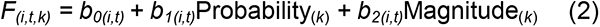

where *F_(i,t,k)_* is the average firing rate of neuron *i* at time t on trial *k,* Probability_(*k*)_ is the probability of reward cued to the monkey in trial *k,* and Magnitude_(*k*)_ *is* the magnitude of reward cued to the monkey in trial *k*. The regression coefficients *b_0(i,t)_* to *b_2(i,t)_* describe the degree to which the firing rates of neuron *i* depend on the mean firing rates (hence, firing rates independent of task parameters), probability of rewards, and magnitude of rewards, respectively, at a given time *t* during the trials.

*Exp. 2.* We first set six items and four locations as categorical parameters. We then described the average firing rates of neuron *i* at time *t* as a linear combination of item and location in each neural population:

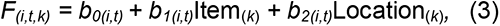

where *F_(i,t,k)_* is the average firing rate of neuron *i* at time t on trial *k,* Item_(*k*)_ is the type of item cued to the monkey on trial *k,* and Location_(*k*)_ *is* the type of location cued to the monkey on trial *k*. Each of the regression coefficients *b_0(i,t)_*, *b_1(i,t)_*, and *b_2(i,t)_* describe the degree to which the firing rates of neuron *i* depend on the mean firing rates (hence, firing rates independent of task parameters, probability, and magnitude of rewards), the degree of the firing rate in each item relative to the mean firing rates, and the degree of firing in each location relative to the mean firing rates, respectively, at a given time *t* during the trials. Note that the interaction term was not included in the model.

We used the regression coefficients (i.e., the regression table in the ANOVA) described in Eqs. 2 and 3 to identify how the dimensions of neural-population signals were composed of information related to probability and magnitude (Exp. 1) and were composed of information related to item and location (Exp. 2) as aggregated properties of individual neural activity. This step constructs an encoding model where the regression coefficients could be explained by a temporal structure in the neural modulation of two continuous parameters (Exp. 1) or two categorical parameters (Exp. 2) at the population level. Our procedures are analogous to the state-space analysis performed by Mante et al. (Mante et al., 2013), in which the regression coefficients were used to provide an axis (or dimension) of the parameters of interest in multi-dimensional state space obtained through principal component analysis (PCA). In this study, our orthogonalized task design allowed us to reliably project the neural firing rates into the regression subspace. Note that our analyses were not aimed at describing the population dynamics of neural signals as a trajectory in multi-dimensional task space but were aimed at describing the neural-modulation dynamics as in a representational model.

#### Preference ordering

In Exp. 2, each neuron had a preferred item and location. As in the conventional representational-model analysis, we defined the preferred item and location in each neuron to construct matrix *X*. We constructed *X* with and without rank order. Items 1 to 6 were rank-ordered from the most preferred to least preferred, defined as the mean firing rates during a whole analysis time window from 0.08 to 0.6 s. Thus, Item_(*k*)_ was the rank-ordered item cued to the monkey on trial *k.* In the same way as the definition of Item, Location_(*k*)_ was the rank-ordered location cued to the monkey on trial *k*. Note that this preference ordering was never changed through time *t* in each neuron *n*.

##### Principal Component Analysis

We used PCA to identify the dimensions of the neural-population signal in the orthogonal spaces composed of the probability and magnitude of rewards in Exp. 1 and of the item and location in Exp. 2, respectively, in each of the four neural populations. In each neural population, we first prepared a two-dimensional data matrix *X* of size *N_(n)_*×*M _(C_*_×*T)*_; the regression coefficient vectors, *b_1(i,t)_* and *b_2(i,t)_*, in Eqs. 2 and 3, whose rows correspond to the total number of neurons (*n*) in each neural population and columns correspond to *C,* the total number of conditions (i.e., two: probability and magnitude in Exp. 1; 10: six items and four locations in Exp. 2), and T as the total number of the analysis windows (i.e., 30 bins: 0.6 s divided by the window size bin, 0.02 s). A series of eigenvectors was obtained by applying PCA once to data matrix *X* in each of the neural populations. The principal components (PCs of this data matrix are vectors *v_(a)_* of length *N_(n)_*, and the total number of recorded neurons if *M _(C_*_×*T)*_ is > *N_(n)_*; otherwise, the length is *M _(C_*_×*T)*_. The PCs were indexed from the principal components, explaining most of the variance to the least variance. The eigenvectors were obtained using the prcomp () function in R software. Note that we did not include the intercept term *b_0(i,t)_* to focus on the neural modulation by the interested parameters.

#### Eigenvectors

When we applied PCA to data matrix *X*, we decomposed the matrix into eigenvectors and eigenvalues. Each eigenvector has a corresponding eigenvalue. In our analysis, the eigenvectors at time *t* represent a vector in the space of probability and magnitude in Exp. 1 and of item and location in Exp. 2, respectively. The eigenvalues at time *t* for the probability and magnitude in Exp. 1 and of item and location in Exp. 2, respectively, were scalars, indicating the extent of variance in the data in that vector. Thus, the first PC is the eigenvector with the highest eigenvalue. We mainly analyzed eigenvectors for the first three PCs (PC1 to PC3) in the following analyses, as the top three PCs had been analyzed previously (Okazawa et al., 2021). Note that we applied PCA once to each neural population, and thus, the total variances contained in the data differed among the neural populations.

#### Analysis of eigenvectors

We evaluated the characteristics of eigenvectors for PC1 to PC3 in each neural population in terms of the vector angle, size, and deviance in the space of probability and magnitude in Exp. 1 and of the item and location in Exp. 2, respectively. The angle is the vector angle from the horizontal axis from 0° to 360° against the main PCs. Size is the length of the eigenvector. The deviance is the difference between vectors. We estimated the deviance from the mean vector for each neural population. These three characteristics of the eigenvectors were compared in each population at *P* < 0.05 using the Kruskal–Wallis and Wilcoxon rank-sum tests. The vector during the first 0.1 s was extracted from these analyses.

#### Shuffle control for PCA

We performed three shuffle controls to examine the significance of population structures described with PCA. A two-dimensional data matrix *X* was randomized by shuffling in three ways. In shuffled control 1, matrix *X* was shuffled by permutating the allocation of neuron *n* at each time *i*. This shuffle provided a data matrix *X* of size *N_(n)_*×*M _(C_*_×*T)*_, eliminating the temporal structure of neural modulation by condition *C* in each neuron but retaining the neural modulations at time *t* at the population level. In shuffled control 2, matrix *X* was shuffled by permutating the allocation of time *i* in each neuron *n*. This shuffle provided a data matrix *X* of size *N_(neuron)_*×*M _(C_*_×*T)*_, eliminating the neural modulation structure under condition *C* maintained in each neuron but retaining the neural modulation in each neuron at the population level. In shuffled control 3, matrix *X* was shuffled by permutating the allocation of both time *i* and neuron *n*. In these three shuffle controls, matrix *X* was estimated 1,000 times. PCA performance was evaluated by constructing the distributions of explained variances for PC1 to PC12. The statistical significance of the variances explained by PC1 and PC3 was estimated based on the 95th percentile of the reconstructed distributions of explained variance or bootstrap standard errors (i.e., standard deviation of the reconstructed distribution).

##### Matrix Size Control for PCA

Because the original matrix sizes of *X*, *N_(n)_*×*M _(C_*_×*T)*_, differed between the cOFC (*X* of size *N_(190)_*×*M _(2_*_×*30*)_) and HPC (*X* of size *N_(590)_*×*M _(10_*_×*30)*_) *populations,* we controlled for matrix size. In this control, we used only two columns in each bin, the most preferred and least preferred, for each condition C, item, and location; thus, matrix X was (*X* of size *N_(590)_*×*M _(4_*_×*30)*_). This corresponds to the conventional analysis usually used in the representational model, which compares the neural responses between the most preferred and least preferred conditions. We evaluated the percentage explained by the model between the original matrix and size-controlled matrix in the HPC.

## Acknowledgments

The authors would like to thank Takashi Kawai, Ryo Tajiri, Yoshiko Yabana, and Yuki Suwa for their technical assistance. The authors would like to thank Tomohiko Takei and Yasuhiro Tsubo for their discussions. Monkey FU was provided by NBRP “Japanese Monkeys” through the National Bio Resource Project of the MEXT, Japan. Funding: This research was supported by JSPS KAKENHI Grant Number JP:15H05374, 19H05007, 21H02797, Takeda Science Foundation, Research Foundation for the Electrotechnology of Chubu (H.Y.), The National Natural Science Foundation of China (Grant 31871139) (Y.N.).

## Author Contributions

H.Y. conceptualized the study. H.Y. and Y.N. designed the experiments. H.Y., H.C., Y.I., Y.H., and T.M. conducted the experiments. M.M. conducted a part of experiments. H.Y. developed the analytic tools. H.Y. and H.C. analyzed the data. H.Y., H.C., J.K., T.O., T.M., and Y.N. evaluated the results. H.Y., H.C., J.K., T.O., and T.M. wrote the manuscript. All authors edited and approved the final manuscript.

## Data availability

All data and analysis codes in this study are available from the corresponding authors.

## References

Aoi, M. C., Mante, V., & Pillow, J. W. (2020). Prefrontal cortex exhibits multidimensional dynamic encoding during decision-making. Nat Neurosci, 23(11), 1410–1420. doi:10.1038/s41593-020-0696-5

Aoi, M. C., & Pillow, J. W. (2018). Model-based targeted dimensionality reduction for neuronal population data. Adv Neural Inf Process Syst, 31, 6690–6699.

Balleine, B. W., Delgado, M. R., & Hikosaka, O. (2007). The role of the dorsal striatum in reward and decision-making. J Neurosci, 27(31), 8161–8165.

Brendel, W., Romo, R., & Machens, C. K. (2011). Demixed Principal Component Analysis. Advances in Neural Information Processing Systems, 24, 2654–2622.

Buzsaki, G., Stark, E., Berenyi, A., Khodagholy, D., Kipke, D. R., Yoon, E., & Wise, K. D. (2015). Tools for probing local circuits: high-density silicon probes combined with optogenetics. Neuron, 86(1), 92–105. doi:10.1016/j.neuron.2015.01.028

Chen, H., & Naya, Y. (2020). Forward Processing of Object-Location Association from the Ventral Stream to Medial Temporal Lobe in Nonhuman Primates. Cereb Cortex, 30(3), 1260–1271. doi:10.1093/cercor/bhz164

Chen, X., & Stuphorn, V. (2015). Sequential selection of economic good and action in medial frontal cortex of macaques during value-based decisions. Elife, 4. doi:10.7554/eLife.09418

Churchland, M. M., Cunningham, J. P., Kaufman, M. T., Foster, J. D., Nuyujukian, P., Ryu, S. I., & Shenoy, K. V. (2012). Neural population dynamics during reaching. Nature, 487(7405), 51–56. doi:10.1038/nature11129

Dayan, P., & Abbott, L. (2001). Theoretical neuroscience: computational and mathematical modeling of neural systems.

Ebitz, R. B., & Hayden, B. Y. (2021). The population doctrine in cognitive neuroscience. Neuron, 109(19), 3055–3068. doi:10.1016/j.neuron.2021.07.011

Elsayed, G. F., & Cunningham, J. P. (2017). Structure in neural population recordings: an expected byproduct of simpler phenomena? Nat Neurosci, 20(9), 1310–1318. doi:10.1038/nn.4617

Evarts, E. V. (1968). A technique for recording activity of subcortical neurons in moving animals. Electroencephalogr Clin Neurophysiol, 24(1), 83–86.

Fetz, E. E., & Cheney, P. D. (1980). Postspike facilitation of forelimb muscle activity by primate corticomotoneuronal cells. J Neurophysiol, 44(4), 751–772. doi:10.1152/jn.1980.44.4.751

Georgopoulos, A. P., Kalaska, J. F., Caminiti, R., & Massey, J. T. (1982). On the relations between the direction of two-dimensional arm movements and cell discharge in primate motor cortex. J Neurosci, 2(11), 1527–1537.

Grafen, A., & Hails, R. (2002). Modern Statistics for the Life Sciences. New York: Oxford university press.

Hubel, D. H., & Wiesel, T. N. (1959). Receptive fields of single neurones in the cat’s striate cortex. J Physiol, 148, 574–591. doi:10.1113/jphysiol.1959.sp006308

Humphries, M. D. (2021). Strong and weak principles of neural dimension reduction. *Neurons, Behavior*, Data analysis, and Theory, 5(2).

Jones, J. P., & Palmer, L. A. (1987). An evaluation of the two-dimensional Gabor filter model of simple receptive fields in cat striate cortex. J Neurophysiol, 58(6), 1233–1258. doi:10.1152/jn.1987.58.6.1233

Jun, J. J., Steinmetz, N. A., Siegle, J. H., Denman, D. J., Bauza, M., Barbarits, B., … Harris, T.D. (2017). Fully integrated silicon probes for high-density recording of neural activity. Nature, 551(7679), 232–236. doi:10.1038/nature24636

Keemink, S. W., & Machens, C. K. (2019). Decoding and encoding (de)mixed population responses. Curr Opin Neurobiol, 58, 112–121. doi:10.1016/j.conb.2019.09.004

Mante, V., Sussillo, D., Shenoy, K. V., & Newsome, W. T. (2013). Context-dependent computation by recurrent dynamics in prefrontal cortex. Nature, 503(7474), 78–84. doi:10.1038/nature12742

Mountcastle, V., & Henneman, E. (1949). Pattern of tactile representation in thalamus of cat. J Neurophysiol, 12(2), 85–100. doi:10.1152/jn.1949.12.2.85

Murray, J. D., Bernacchia, A., Roy, N. A., Constantinidis, C., Romo, R., & Wang, X. J. (2017). Stable population coding for working memory coexists with heterogeneous neural dynamics in prefrontal cortex. Proc Natl Acad Sci U S A, 114(2), 394–399. doi:10.1073/pnas.1619449114

O’Keefe, J., & Dostrovsky, J. (1971). The hippocampus as a spatial map. Preliminary evidence from unit activity in the freely-moving rat. Brain Res, 34(1), 171–175. doi:10.1016/0006-8993(71)90358-1

Okazawa, G., Hatch, C. E., Mancoo, A., Machens, C. K., & Kiani, R. (2021). Representational geometry of perceptual decisions in the monkey parietal cortex. Cell, 184(14), 3748–3761 e3718. doi:10.1016/j.cell.2021.05.022

Osako, Y., Ohnuki, T., Tanisumi, Y., Shiotani, K., Manabe, H., Sakurai, Y., & Hirokawa, J. (2021). Contribution of non-sensory neurons in visual cortical areas to visually guided decisions in the rat. Curr Biol, 31(13), 2757–2769 e2756. doi:10.1016/j.cub.2021.03.099

Platt, M. L., & Glimcher, P. W. (1999). Neural correlates of decision variables in parietal cortex. Nature, 400(6741), 233–238.

Raposo, D., Kaufman, M. T., & Churchland, A. K. (2014). A category-free neural population supports evolving demands during decision-making. Nat Neurosci, 17(12), 1784–1792. doi:10.1038/nn.3865

Rossi-Pool, R., Zainos, A., Alvarez, M., Diaz-deLeon, G., & Romo, R. (2021). A continuum of invariant sensory and behavioral-context perceptual coding in secondary somatosensory cortex. Nat Commun, 12(1), 2000. doi:10.1038/s41467-021-22321-x

Saxena, S., & Cunningham, J. P. (2019). Towards the neural population doctrine. Curr Opin Neurobiol, 55, 103–111. doi:10.1016/j.conb.2019.02.002

Tolhurst, D. J., & Movshon, J. A. (1975). Spatial and temporal contrast sensitivity of striate cortical neurones. Nature, 257(5528), 674–675. doi:10.1038/257674a0

Vyas, S., Golub, M. D., Sussillo, D., & Shenoy, K. V. (2020). Computation Through Neural Population Dynamics. Annu Rev Neurosci, 43, 249–275. doi:10.1146/annurev-neuro-092619-094115

Wurtz, R. H. (1968). Visual cortex neurons: response to stimuli during rapid eye movements. Science, 162(3858), 1148–1150. doi:10.1126/science.162.3858.1148

Yamada, H., Imaizumi, Y., & Matsumoto, M. (2021). Neural Population Dynamics Underlying Expected Value Computation. J Neurosci, 41(8), 1684–1698. doi:10.1523/JNEUROSCI.1987-20.2020

Yamada, H., Inokawa, H., Matsumoto, N., Ueda, Y., Enomoto, K., & Kimura, M. (2013). Coding of the long-term value of multiple future rewards in the primate striatum. J Neurophysiol, 109(4), 1140–1151. doi:10.1152/jn.00289.2012

Yamada, H., Louie, K., Tymula, A., & Glimcher, P. W. (2018). Free choice shapes normalized value signals in medial orbitofrontal cortex. Nat Commun, 9(1), 162. doi:10.1038/s41467-017-02614-w

Yuste, R. (2015). From the neuron doctrine to neural networks. Nat Rev Neurosci, 16(8), 487–497. doi:10.1038/nrn3962

